# Stability of influenza A H5N1 virus in raw milk cheese

**DOI:** 10.1101/2025.03.13.643009

**Authors:** Mohammed Nooruzzaman, Pablo Sebastian Britto de Oliveira, Nicole H. Martin, Samuel D. Alcaine, Diego G. Diel

**Author notes:** Corresponding author: Diego G. Diel;.

## Abstract

Here we evaluated the stability of highly pathogenic avian influenza (HPAI) H5N1 virus in raw milk cheeses using a mini cheese model prepared with HPAI-spiked raw milk under varying pH levels (pH 6.6, 5.8 and 5.0) and in commercial raw milk cheese inadvertently produced with naturally contaminated raw milk. We observed a pH-dependent survival of the virus, with infectious virus persisting throughout the cheese making process and for up to 60 days of aging in the pH 6.6 and 5.8 cheese groups. Whereas at pH 5.0, the virus did not survive the cheese making process. These findings were validated using the commercial raw milk cheese samples in which infectious virus was detected for up to 60 days of aging. Our study highlights the potential public health risks of consuming raw milk cheese, underscoring the need for additional mitigation steps in cheese production to prevent human exposure to infectious virus.

## Main text

The spillover of highly pathogenic avian influenza (HPAI) H5N1 virus clade 2.3.4.4b genotype B3.13 and more recently of genotype D1.1 into dairy cows ^1,2^ and their continuous circulation in cattle in the United States^3^ poses major animal and public health concerns. The tropism of the virus for the mammary gland milk secreting epithelial cells, leading to severe viral mastitis^1,4,5^ and shedding of high levels of infectious virus in milk (up to 8.8 log_10_ TCID_50_/mL), pose a risk of exposure to other animals (including dairy cows, pets, birds, and wildlife) and potentially humans^1,6–9^. Although recent studies have shown that pasteurization and various thermization conditions effectively inactivate HPAI H5N1 virus in milk^10–13^, consumption of raw milk and raw milk cheeses, which are not subjected to thermal treatment, remains popular worldwide, including in the United States and represents a unique exposure risk to infectious virus.

A survey conducted between 2016 and 2019 in the U.S. by FDA showed that 4.4% of U.S adults reported consuming raw milk at least once a year, with 1.6% consuming it frequently^14^ and 1.6% of the population consuming raw milk cheeses^15^. These findings illustrate the high risk to public health, as HPAI H5N1 virus has been shown to persist in raw milk under refrigeration (4°C) for at least 8 weeks^11,12^ and many states in which the virus has been detected in dairy cattle allow commercialization of raw milk dairy products, although no oral infection from raw milk ingestion has been documented to date in humans. Importantly, in California, one of the states with the largest number of herds affected by HPAI H5N1 virus^3^, voluntary recalls of several raw milk dairy products were issued in November and December 2024 after retail raw milk tested positive for H5N1^16^.

To enhance the safety of raw milk cheeses, the U.S. Code of Federal Regulation (21 CFR Part 133 – Cheeses and Related Cheese Products) requires that cheeses produced from unpasteurized (raw) milk must undergo a curing process for a minimum of 60 days at temperatures no lower than 35 °F (1.67°C) to inactivate bacterial pathogens. However, research has not yet determined whether the HPAI H5N1 virus is inactivated through the intricate cheese making process and the mandatory aging period. ^14^

In this study we investigated the survival of HPAI H5N1 virus in raw milk cheeses using two approaches. First, we developed a mini-cheese model to examine viral stability during cheese production and aging under varying pH conditions, using raw milk spiked with H5N1 virus. Additionally, we analyzed viral stability in commercial raw milk cheeses inadvertently made with naturally contaminated milk.

To evaluate the effect of pH on HPAI H5N1 virus stability using the raw milk mini-cheese model, we selected three pH levels (6.6, 5.8, and 5.0), with these values being chosen based on a survey of 273 commercial cheese samples used to established a categorization framework for critical physicochemical parameters of cheeses^17^. Cheeses were made by direct acidification (lactic acid) of raw milk spiked with HPAI H5N1 TX2/24 virus (clade 2.3.4.4b genotype B3.13) (**Extended Data Fig. 1**). Milk, whey and curd samples were collected during cheese production to quantify viral RNA (real-time reverse transcriptase PCR [rRT-PCR]) and infectious virus (titrations in embryonated chicken eggs [ECEs]). Viral RNA loads in milk before (7.0 ± 0.42-7.16 ± 0.09 log_10_ copy number/mL) and after (6.6 ± 0.51-7.04 ± 0.19 log_10_ copy number/mL) pH adjustment and after heat treatment at 34°C for 1h (6.78 ± 0.22-7.01 ± 0.3 log_10_ copy number/mL) were comparable in all three pH conditions but were slightly lower (4.93 ± 0.66-5.96 ± 0.22 log_10_ copy number/mL) in whey (**Fig. 1a**). Notably, lower viral RNA loads (4.93 ± 0.66-5.85 ± 0.14 log_10_ copy number/mL) were detected in whey and curd samples in the pH 5.0 cheese group when compared to corresponding samples from the pH 6.6 (5.88 ± 0.28-6.38 ± 0.33 log_10_ copy number/mL) and 5.8 (5.87 ± 0.19-6.23 ± 0.02 log_10_ copy number/mL) groups (**Fig. 1a**).

**Fig. 1.**
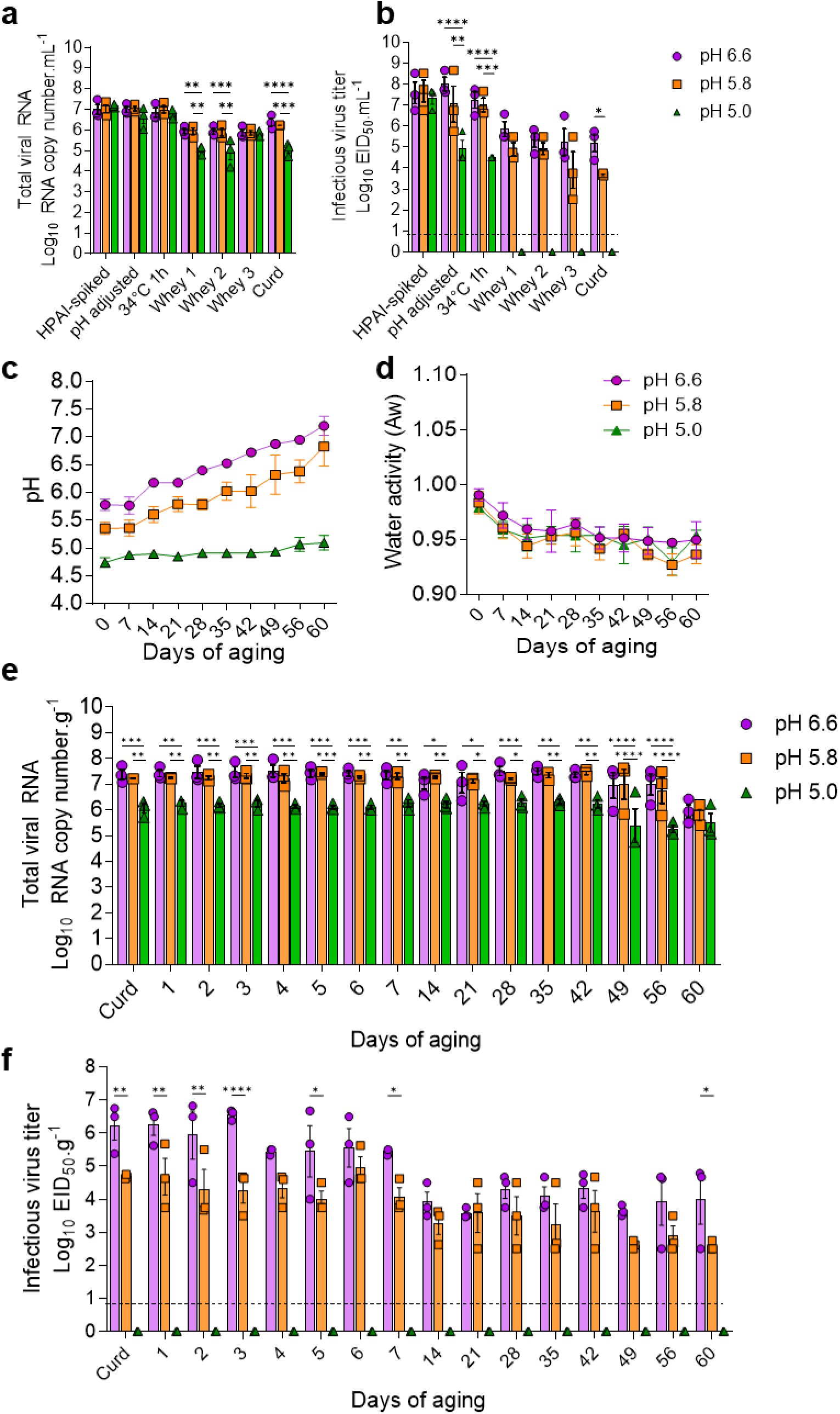
Long-term stability of HPAI H5N1 virus in raw milk cheese. HPAI H5N1 viral RNA (**a**) and infectious virus (**b**) loads in HPAI-spiked raw milk and whey during cheese making process in cheeses made with milk at pH 6.6, 5.8, and 5.0 as determined by rRT-PCR and virus titration in embryonated chicken eggs (ECEs), respectively (n = 3). Changes in pH (**c**) and water activity (**d**) of raw milk cheese during a 60-day aging period at 4°C. HPAI H5N1 viral RNA (**e**) and infectious virus (**f**) loads in raw milk cheese during the 60-day aging process as determined by rRT-PCR and virus titration in embryonated chicken eggs, respectively (n = 3). 2-way ANOVA followed by multiple comparisons test, * *p* < 0.05, ** *p* < 0.01, *** *p* < 0.001 and **** *p* < 0.0001. The results represent the mean ± standard error of three independent experiments.

Infectious virus titers in HPAI H5N1 virus-spiked raw milk ranged between 7.34 ± 0.51-7.68 ± 0.89 log_10_ EID_50_/mL (**Fig. 1b**). After adjustment of the milk pH at setting, infectious virus levels remained stable in the pH 6.6 (8.02 ± 0.57 log_10_ EID_50_/mL) and 5.8 (7.06 ± 1.48 log_10_ EID_50_/mL) cheese groups, whereas a significant decrease in infectious virus titers (4.5 ± 0.0-4.92 ± 0.72 log_10_ EID_50_/mL; p<0.01) was observed in the pH 5.0 cheese group on samples collected after pH adjustment and after incubation at 34°C for 1 h (**Fig. 1b**). Infectious virus was also detected in whey samples (3.68 ± 0.06-5.86 ± 0.63 log_10_ EID_50_/mL) collected from the pH 6.6 and 5.8 cheese groups after coagulation (whey 1), salting (whey 2), and whey draining (whey 3), and in the cheese curd (**Fig. 1b**). Viral titers were slightly lower (3.68 ± 0.06-4.93 ± 0.49 log_10_ EID_50_/mL) in whey 1-3 and curd samples collected from the pH 5.8 cheese group when compared to pH 6.6 (5.21 ± 0.73-5.86 ± 0.63 log_10_ EID_50_/mL) (**Fig. 1b**). Notably, no infectious virus was detected in whey and curd samples collected from pH 5.0 cheese group (**Fig. 1b**). These results demonstrate a pH-dependent stability of HPAI H5N1 virus during raw milk cheese production

Next, we evaluated the stability of HPAI H5N1 virus during cheese aging or curing at 4°C over a 60-day period. Cheese samples were collected daily between days 1 and 7 and then on days 14, 21, 35, 42, 49, 56, and 60 of aging and used to monitor the cheese pH, water activity (Aw) and viral stability in the cheese matrix over time. The mean pH of the curd in the pH 6.6, 5.8 and 5.0 cheese groups were significantly lower (5.78 ± 0.18; 5.36 ± 0.2; and 4.74 ± 0.15, respectively) than the milk pH at setting (**Fig. 1c**). During the first week of aging, the mean pH of the 6.6 and 5.8 cheese groups remained constant, but they gradually increased (by ∼1.4 units) in the subsequent weeks until the end of the 60-day aging period (**Fig. 1c**). In contrast, the mean pH of the pH 5.0 cheese group was relatively stable, only slightly increasing by 0.35 units from 4.74 ± 0.15 to 5.09 ± 0.23 over the 60-day aging period (**Fig. 1c)**. The mean water activity (A_w_) in the curd of the pH 6.6, 5.8 and 5.0 cheese groups were comparable (0.990 ± 0.01, 0.983 ± 0.017, and 0.979 ± 0.005, respectively) (**Fig. 1d**). The A_w_ in all three cheese groups markedly decreased in the first 14 days of aging (**Fig. 1d**). This trend continued, though at a slower rate, until day 60 of aging (**Fig 1d**).

The HPAI H5N1 viral RNA remained stable in raw milk cheeses throughout the entire 60-day aging period (**Fig. 1e**). While viral RNA levels were comparable in the pH 6.6 and 5.8 cheese groups, approximately a 1-log reduction in viral RNA loads was observed in the pH 5.0 cheese group (**Fig. 1e**). Consistently high amounts of infectious virus were recovered from raw milk cheeses in the pH 6.6 and 5.8 cheese groups during the first 7 days of aging (5.81 ± 0.64 and 4.38 ± 0.67 log_10_ EID_50_/g, respectively) (**Fig. 1f**). Notably, virus titers only dropped approximately 2 logs over the next 60 days of aging, resulting in viral yields of 3.99 ± 1.29 and 2.58 ± 0.14 log_10_ EID_50_/g in the pH 6.6 and 5.8 cheese groups, respectively (**Fig. 1f**). Of note, the viral titers recovered in the pH 5.8 cheese group were 1.5-2.0 logs lower than in the pH 6.6 cheese group throughout the 60-day aging period. Importantly, no infectious virus was recovered from raw milk cheeses at the pH 5.0 group (**Fig. 1f**). Virus infectivity in allantoic fluid of ECEs inoculated with the cheese homogenates was confirmed by hemagglutination assay (**Supplementary Data Table 1**) throughout the 60-day aging period and by rRT-PCR targeting the influenza M gene on day 60 samples (**Extended Data Fig. 2**). The decimal reduction times, *D*-values, of H5N1 virus in the raw milk cheeses in the pH 6.6 and 5.8 groups were estimated to be 25.5 and 32.2 days, respectively, indicating long-term stability of the virus in raw milk cheese.

We also investigated the HPAI H5N1 virus stability in commercial raw milk cheeses from a raw milk dairy that inadvertently made cheddar cheese with HPAI H5N1 contaminated raw milk, following an H5N1 influenza virus outbreak in the farm. We received four (n = 4) commercial raw milk cheese blooks (2 pounds/block) on day 24 of aging. Upon arrival, these cheese samples were tested for the presence of HPAI H5N1 virus by rRT-PCR and virus titrations in ECEs and the pH (5.37 ± 0.06) and A_w_ (0.9416 ± 0.005) on each sample was recorded. Following testing and confirmation that the cheese blocks were indeed positive for H5N1 virus genotype B3.13 (**Extended Data Fig. 3**) we continued the aging process at 4°C for 60 days. Individual samples (1 g) were collected from each commercial raw milk cheese block on days 29, 32, 35, 38, 42, 45, 49, 52, 56 and 60 of aging. Additionally, the pH and A_w_ of the cheeses were recorded at the same point. The mean pH of the commercial cheeses, which initiated at 5.37 ± 0.06 on day 24, remained relatively stable until day 60 of aging (5.34 ± 0.2) (**Fig. 2a**). As expected, there was a slight reduction in A_w_ during the aging process, which initiated at 0.9416 ± 0.005 on day 24 and was determined to be 0.9284 ± 0.011 on day 60 of aging (**Fig. 2b**). The rRT-PCR analysis revealed the presence of high viral RNA loads (5.82 ± 0.37 log10 copy number/g) in the commercial cheese samples on day 24 which remained stable throughout the 60-day aging period (**Fig. 2c**). Similarly, virus titrations in ECEs showed mean viral titers of 4.21 ± 0.48 log_10_ EID_50_/g of infectious virus in the commercial cheeses on day 24, with slight variations in the viral loads being observed throughout the aging process. Notably, on day 60 of aging 4.0 ± 0.58 log_10_ EID_50_/g of H5N1 virus were recovered from the commercial raw milk cheese samples (**Fig. 2d**). The presence of HPAI H5N1 virus in the allantoic fluids collected from ECEs inoculated with the cheese homogenates was confirmed by HA (**Supplementary Data Table 1**) and rRT-PCR (**Extended Data Fig. 4**). These findings confirm and validate the data obtained in our laboratory-scale mini-cheese model. Most importantly, these results provide compelling evidence that HPAI H5N1 virus is stable in raw milk cheese, surviving throughout the minimum required aging period (60 days) for raw milk cheeses.

**Fig. 2.**
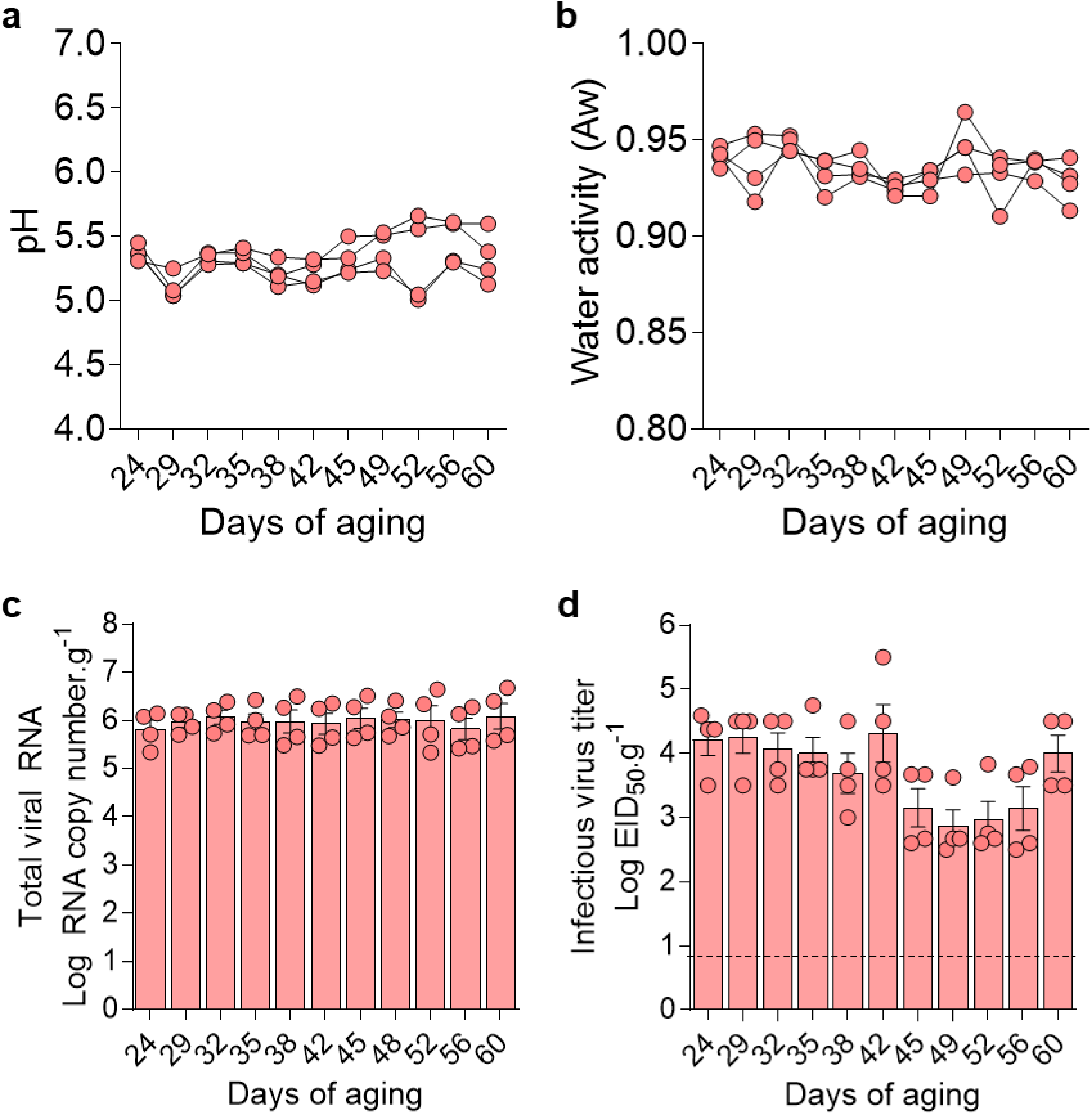
Stability of HPAI H5N1 virus stability in commercial raw milk cheeses. Changes in pH (a) and water activity (b) of commercial raw milk cheese samples during 60-day of aging process. (c) HPAI H5N1 viral RNA loads in commercial raw milk cheese samples during 60-day aging period as determined by rRT-PCR (n = 4). (d) HPAI H5N1 infectious virus loads in commercial raw milk cheese samples during 60-day aging period as determined by virus titration in embryonated chicken eggs (n = 4). The data is presented as individual cheese read outs (a, b, c, d) and as the mean ± standard error (c and d) of four commercial raw milk cheese samples.

Our study demonstrates that HPAI H5N1 virus exhibits remarkable stability throughout the cheese making process, with slow decay rates (*D*-values 25-32 d) and detectable infectious virus persisting for up to 60 days of aging. Notably, we found that pre-processing milk acidification to pH 5.0 (with lactic acid) effectively inactivated the virus, leading to no infectious virus detection in cheese curd immediately after cheese production and subsequently in raw milk cheese throughout the 60-day aging. Furthermore, our recent findings indicate that thermization (sub-pasteurization) of raw milk at temperatures above 54°C successfully inactivates the virus within 15 minutes ^12^, offering an alternative safety measure for cheese production.

Influenza A viruses (IAV) are sensitive to acidic environments similar to those found in cellular endosomes (37°C, pH ∼5.5)^18^. This sensitivity is primarily driven by pH-dependent conformational changes in the viral hemagglutinin (HA) protein^18,19^. Under normal infection conditions, HA transitions to its irreversible post-fusion conformation within the endosome, facilitating the crucial membrane fusion between the viral envelope and the endosomal membrane, thereby enabling viral uncoating and cellular entry^18^. However, if this conformational change occurs prematurely outside the cellular environment, the HA protein loses its ability to bind to host cell receptors, resulting in viral inactivation. This mechanism makes IAV particularly vulnerable to acidic conditions in the external environment. Our results from the raw milk cheese group subjected to pre-processing milk acidification to pH 5.0 provide strong support for this mechanism. Furthermore, using a nano-luciferase reporter H5N1 virus (rTX2/24-luc) in viral entry assays, we demonstrated that acidifying milk to pH 5.5 or lower significantly impairs viral entry into bovine uterine epithelial cells (**Extended Data Fig. 5**), confirming the role of pH in viral inactivation, resulting in impaired viral entry and infectivity.

Our study has implications for public health, food safety, and regulatory policies. First, the current regulation requiring 60-day aging of raw milk cheese before marketing proves insufficient to achieve HPAI H5N1 virus inactivation and guarantee cheese safety. This concern may extend to other raw milk products as the virus can persist for up to 56 days in raw milk under refrigeration^12^. Importantly, consumption of contaminated commercial raw milk has been linked to H5N1 virus infections in domestic cats (Frye et al., personal communication). Although the infectious dose of the virus to humans is not known, ingestion of contaminated raw dairy products repeatedly may increase the probability of infections. Given the continuous circulation and increasing frequency of HPAI H5N1 virus outbreaks in U.S. dairy herds and the recent emergence of the new potentially more virulent virus genotype D.1.1 in dairy cattle^20^, implementing additional mitigation steps such as testing of raw milk bulk tanks or using milk pasteurization, thermization or acidification before cheese making becomes crucial to ensure food safety.

## Acknowledgements

The work was funded by the U.S. Food and Drug Administration (award no. 1U18FD008488-01) and by the New York State Department of Agriculture and Markets (award no. CM04068HM). The authors would like to thank the Cornell EH&S and Biosafety teams for their help in setting up the protocols and procedures to conduct work with HPAI H5N1 virus at the Cornell BSL-3 facilities and the Cornell Teaching Dairy for providing the raw milk samples. The authors would further like to acknowledge the efforts of Timothy Arnold DeMarsh with the cheese making process and Salman L. Butt with the phylogenetic analysis.

## Data availability

All data pertaining to this study are presented in the paper, in the Extended Data and are available from the corresponding author upon request. Source data are provided with this paper.

## Author contributions

Conceptualization: MN, NHM, SDA, DGD; Methodology: MN, PSBO, NHM, SDA, DGD; Software: MN; Validation: MN, PSBO, DGD; Formal analysis: MN; Investigation: MN, PSBO, DGD; Resources: NHM, SDA, DGD; Data Curation: MN; Writing - Original Draft: MN, DGD; Writing - Review & Editing: MN, PSBO, NHM, SDA, DGD; Visualization: MN; Supervision: DGD; Project administration: DGD, Funding acquisition: NHM, SDA, DGD.

## Extended Data Figures

**Extended Data Fig. 1.**
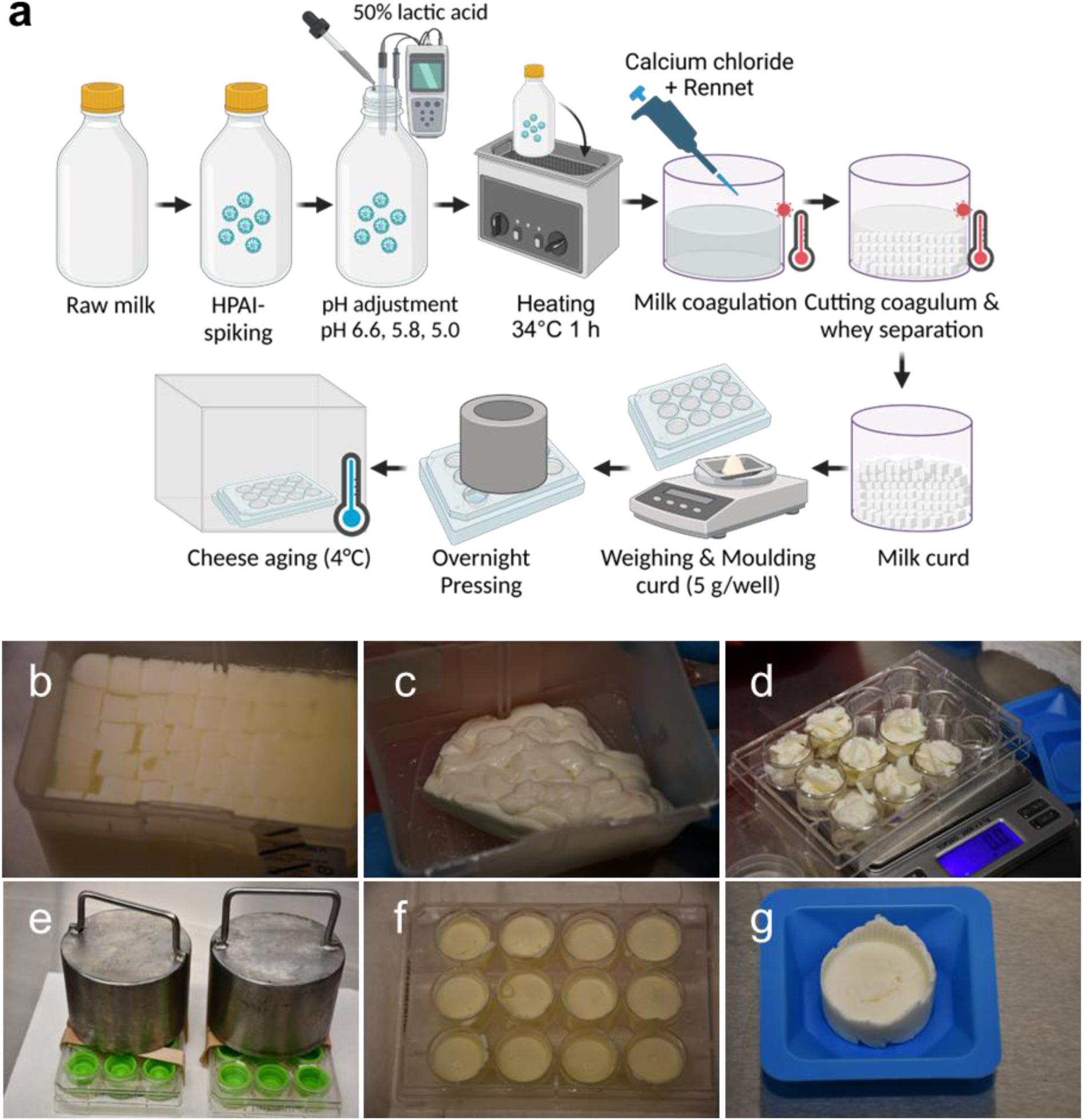
The cheese making process using mini-cheese model. (a) Illustration showing the cheese making process using the mini cheese model developed in this study. Raw normal milk was spiked with HPAI H5N1 and adjusted to different pH conditions with 50% lactic acid. Milk was then pre-warmed at 34°C for 1 h before adding rennet and calcium chloride for milk coagulation at 35°C for 65 min. The milk coagulum was cut using sterile knife and whey was released by applying increment heating (35-43°C) and adding sodium chloride solution. The curd was filtered with sterile cheese cloth for 1 h before aliquoting (5 g/well) into 12-well plate for overnight compression. The compressed cheese samples were aged at 4°C. Photographs during cheese making process showing (b) cut coagulum, (c) curd after whey separation, (d) aliquoting curd into 12-well plate, (e) compression by applying weight, (f-g) compressed cheese blocks.

**Extended Data Fig. 2:**
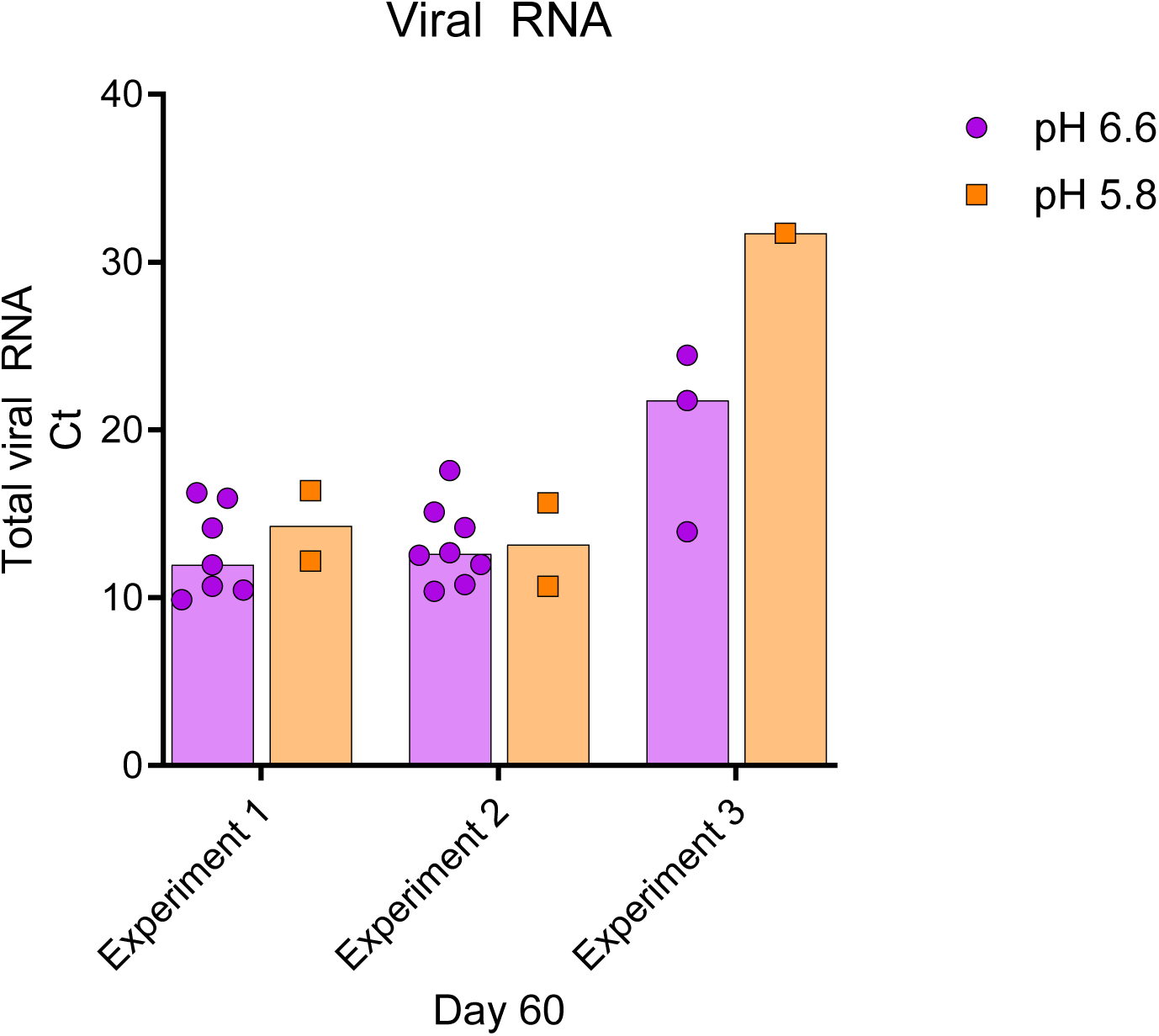
Viral RNA in allantoic fluids of eggs inoculated with day 60 mini cheese samples. Viral RNA loads were quantified by influenza A virus M-gene specific rRT-PCR. Experiment 1-3 are three independent experiments conducted with raw milk cheeses. Data points on each experiment represent the ECEs that were HA positive.

**Extended Data Fig. 3.**
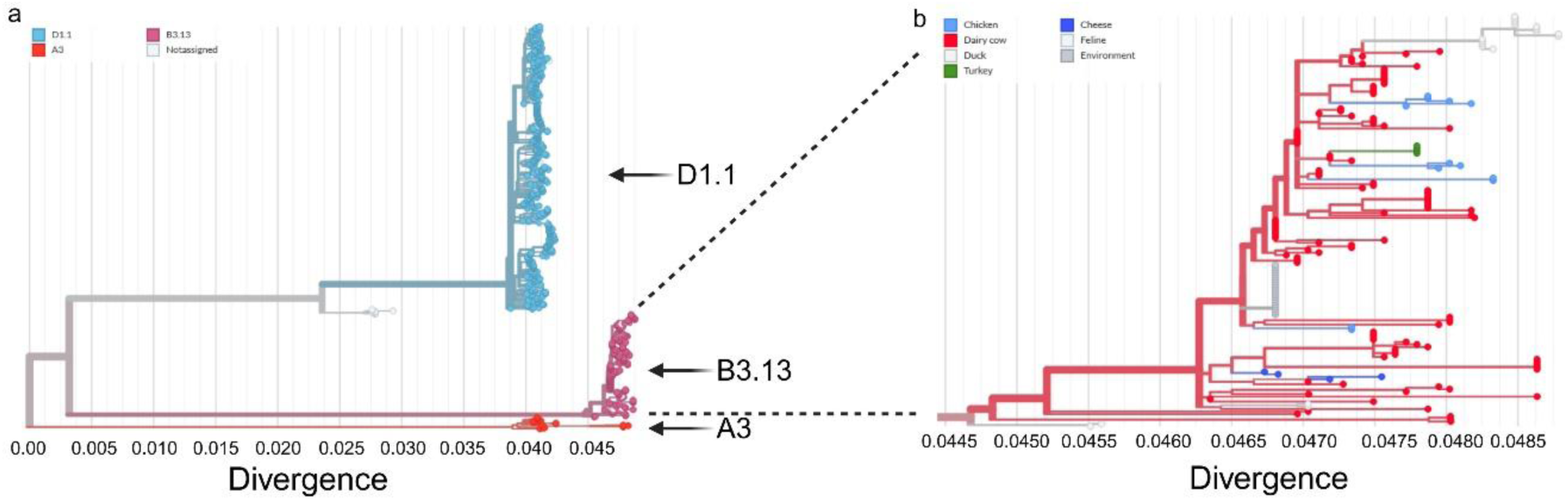
a) Genetic divergence of genotype A3, B3.13, and D1.1 from different hosts. Genetic divergence of B3.13 from four (04) cheese samples and other avian and mammalian hosts.

**Extended Data Fig. 4.**
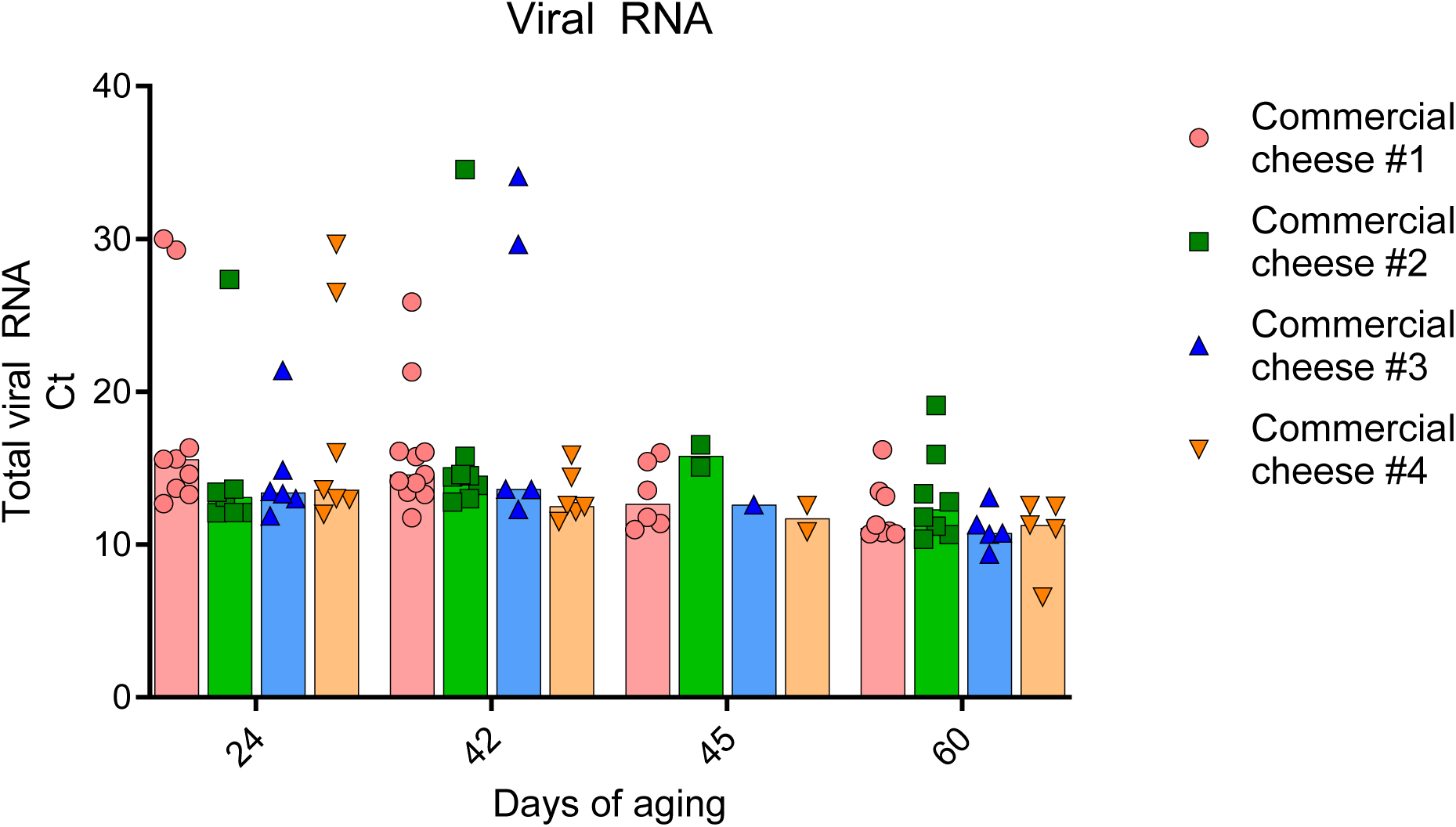
Viral RNA in allantoic fluids of eggs inoculated with commercial cheese samples. Viral RNA loads were quantified by influenza A virus M-gene specific rRT-PCR. Data points on each experiment represent the ECEs that were HA positive.

**Extended Data Fig. 5.**
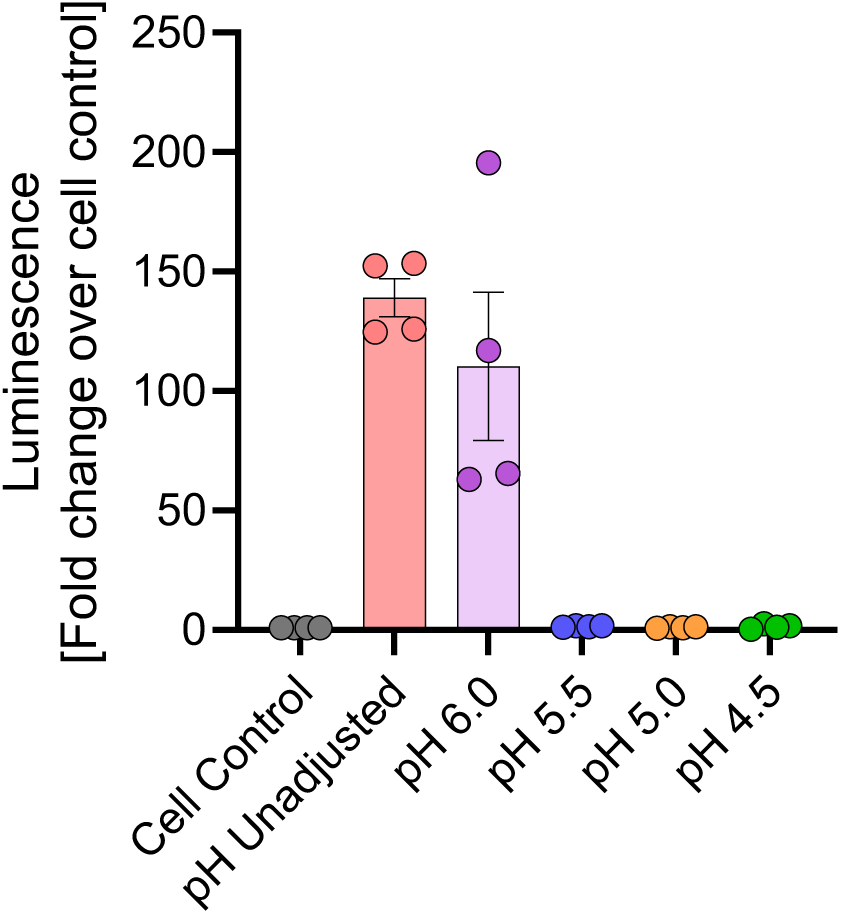
pH dependent infectivity inhibition of HPAI H5N1 in raw milk. Raw normal milk spiked with HPAI H5N1 virus was adjusted to pH 6.0, 5.5, 5.0 and 4.5 and incubated at 4°C for 1 h. Cal-1 cells were inoculated with 0.2 mL milk samples and inoculated at 4°C for 1 h (virus adsorption) and then transferred and incubated at 37°C for 4 h. Cell lysates were harvested, and luciferase activity was measured using a luminometer. The luciferase activity was expressed as fold change over cell control. Data represents observations (dots) from four replicates (n=4) overlayed with the mean (horizontal line) and ± SEM (whiskers) from 2 independent experiments.

**Supplementary Data Table 1.** The hemagglutination (HA) activity of allantoic fluids collected from embryonated chicken eggs (ECEs) inoculated with raw milk cheese samples prepared using mini-cheese model.

|  | **pH 6.6** |  |  | **pH 5.8** |  |  | **pH 5.0** |  |  |
| --- | --- | --- | --- | --- | --- | --- | --- | --- | --- |
| Days | Exp 1 | Exp 2 | Exp 3 | Exp 1 | Exp 2 | Exp 3 | Exp 1 | Exp 2 | Exp 3 |
| 0 | + | + | + | + | + | + | - | - | - |
| 1 | + | + | + | + | + | + | - | - | - |
| 2 | + | + | + | + | + | + | - | - | - |
| 3 | + | + | + | + | + | + | - | - | - |
| 4 | + | + | + | + | + | + | - | - | - |
| 5 | + | + | + | + | + | + | - | - | - |
| 6 | + | + | + | + | + | + | - | - | - |
| 7 | + | + | + | + | + | + | - | - | - |
| 14 | + | + | + | + | + | + | - | - | - |
| 21 | + | + | + | + | + | + | - | - | - |
| 28 | + | + | + | + | + | + | - | - | - |
| 35 | + | + | + | + | + | + | - | - | - |
| 42 | + | + | + | + | + | + | - | - | - |
| 49 | + | + | + | + | + | + | - | - | - |
| 56 | + | + | + | + | + | + | - | - | - |
| 60 | + | + | + | + | + | + | - | - | - |

Note: +, HA positive, -, HA negative

**Supplementary Data Table 2.** Compositional analysis of raw milk used for cheese making in the mini-cheese model.

| **Parameters** | **Batch 1** | **Batch 2** | **Batch 3** | **Batch 4** |
| --- | --- | --- | --- | --- |
| Bacto Standard Plate Count x1000 | 121 | 1 | 1 | 1 |
| Somatic Cell Count x1000 | 93 | 110 | 150 | 210 |
| Butterfat | 4.37 | 4.83 | 4.58 | 4.85 |
| Protein | 3.46 | 3.44 | 3.48 | 3.42 |
| Lactose | 4.99 | 5.02 | 4.98 | 4.99 |
| Other Solids | 5.91 | 5.91 | 5.89 | 5.94 |
| Total Solids | 13.74 | 14.18 | 13.95 | 14.21 |
| Freezing point depression | -0.554 | -0.556 | -0.549 | -0.556 |

## Methods

### Cells

Human kidney cells HEK293T (ATCC CRL-3216) and bovine uterine epithelial cells (Cal-1, developed in house at the Virology Laboratory at the Cornell Animal Health Diagnostic Center, AHDC) were cultured in Modified Eagle Medium (MEM) supplemented with 1% L-glutamine and 10% Fetal Bovine Serum (FBS) and containing penicillin–streptomycin (Thermo Fisher Scientific; 10 U ml–1 and 100 µg ml–1, respectively) at 37°C with 5% CO_2_.

### Virus

The HPAI H5N1 virus isolate TX2/24 (A/Cattle/Texas/063224-24-1/2024, genotype B3.13, GISAID accession number: EPI_ISL_19155861) obtained from pooled milk samples of H5N1 virus infected dairy cows in Texas, USA^1^ was used to spike the raw milk (target titer 10^7^ EID_50_/mL) used for the raw milk cheese studies. The virus stock (passage 3) was propagated in 10-day old embryonated chicken eggs and titrated in embryonated chicken eggs (EID_50_) and Cal-1 cells (TCID_50_). Virus stocks were subjected to whole genome sequencing to determine the integrity of the H5N1 TX2/24 virus sequences. The recombinant HPAI H5N1 TX2/24 expressing NanoLuc luciferase reporter (rTX2/24-NanoLuc) virus was generated in Diel’s Lab using a reverse genetics system as described below.

### Generation of recombinant HPAI TX2/24-NanoLuc reporter virus

A reverse genetics system for the bovine H5N1 virus based on an isolate A/Cattle/Texas/06322424-1/2024 (TX2/24) obtained from milk from infected dairy cows^1^ was established in our laboratory and used as backbone to generate a recombinant virus expressing the Nanoluc luciferase reporter gene (rTX2/24-NanoLuc). Briefly, full length genome sequences of PB1, PB2, PA, HA, NA, NP and M gene segments of TX2/24 strain (H5N1 clade 2.3.4.4b, genotype B3.13, GISAID accession number: EPI_ISL_19155861) were synthesized commercially (Twist Bioscience) and cloned into the dual promoter influenza reverse genetics plasmid pHW2000 (kindly provided by Dr. Richard Webby at St. Jude Children’s Research Hospital) using the BsmBI (New England Biolabs) restriction sites. To generate the NLuc reporter virus, the NS segment of the rTX2/24 recombinant virus was modified to encode a fusion protein (NS-NLuc) from a single nonoverlapping transcript. The NLuc was cloned at the C-terminal of NS1. The NS1 and NEP open reading frames were separated by the porcine teschovirus 1 2A autoproteolytic cleavage site. The NS-NLuc gene segment was synthesized (Twist Bioscience) and cloned into pHW2000 vector using the BsmBI sites. The pHW2000 plasmids containing seven TX2/24 gene segments (PB1, PB2, PA, HA, NA, NP and M) and the modified NS segment encoding NLuc were co-transfected into a co-culture of HEK293T and Cal-1 (bovine uterine epithelial cells) using Lipofectamine 3000 reagents (ThermoFisher Scientific). Cell culture supernatant was harvested after 96 hours and used to infect newly seeded Cal-1 cells. Both cell lysate and culture supernatant were harvested after 72-96 hours to prepare the seed stock for the rTX2/24-NLuc virus. The working stock of the virus was prepared after inoculating 10-day old embryonated chicken eggs via the allantoic cavity route and the infected allantoic fluid was harvested after 48 hours. Viruses from the initial rescue and from passages 1 and 2 were sequenced to confirm the integrity of the sequences and absence of unwanted mutations. The 50% tissue culture infectious dose (TCID_50_) was determined using end-point dilutions and the Spearman and Karber’s method and expressed as TCID_50_.mL^−1^. The sequenced verified stock rTX2/24-NLuc virus was used in the pH dependent inhibition of HPAI H5N1 virus assay.

### Biosafety and biosecurity

All work involving handling and propagation of HPAI H5N1 virus, cheese making and processing, and virus isolation in embryonated chicken eggs were performed following strict biosafety measures in the Animal Health Diagnostic Center (AHDC) research BSL-3 laboratories at the College of Veterinary Medicine, Cornell University.

### Milk samples

Raw normal bulk tank milk used for making cheese in our BSL-3 laboratory was obtained from the Teaching Dairy at the College of Veterinary Medicine at Cornell University. The physical, chemical and microbiological properties and parameters of the milk samples were analyzed prior to cheese making in a third-party laboratory (DairyOne, Ithaca, NY) (**Supplementary Data Table 2**).

### Development of a mini cheese model and production of raw milk mini cheese

To assess the stability of HPAI H5N1 virus in raw milk cheese, our team developed a 5 g mini cheese model and implemented this model in our BSL3 laboratory. For this, 600 mL of raw bulk tank milk were placed in a sterile glass bottle and spiked with the HPAI H5N1 virus isolate TX2/24 virus to obtain a target titer of about 10^7^ EID_50_ or 10^6^ TCID50 per mL of milk. Next, the pH of the HPAI H5N1 virus-spiked milk was measured using a pH meter (HI2002-01 edge® Dedicated pH, Scientific Equipment Source, ON, Canada) and then adjusted to three different conditions, pH 6.6, 5.8 and 5.0 using 50% lactic acid (Sigma Aldrich, USA) to investigate the effect of pH in HPAI H5N1 virus stability in cheese. Following pH adjustment, the milk was warmed at 34°C for 1 h in a water bath. Samples (1 mL aliquots) of raw milk were collected after warming and stored at -80°C for virus titrations. After incubation we confirmed that the milk had reached 32°C and then we proceeded with the coagulation step, by adding 1.875 mL of 32% calcium chloride (GetCulture, WI, USA), and 78 µL of rennet (Chy-Max, Chr. Hansen A/S, Denmark). The milk bottle was then inverted 25 times for mixing and then transferred into a sterile empty pipette tip (1250 µl) box. The box was placed into a zip lock bag, sealed, and then incubated in a water bath at 35°C for 65 min for coagulum formation. After incubation we confirmed that the milk temperature reached 35°C. The coagulated curd mass (coagulum) was then cut with a sanitized knife, and the tip box was returned to the water bath at 35°C for 10 min to heal the coagulum and release the whey from the curd. The coagulum and whey were then heated gradually over the course of 30 min in two phases until it reached 43°C. In the first phase, the temperature of the water bath was increased from 35°C to 39°C over a 15 min period (1 degree increase at every 2 min, then the temperature was held at 39°C for the remainder of 15 min). Similarly, in the second phase, the temperature of the water bath was increased from 39°C to 43°C over a 15 min period (1 degree increase at every 2 min, then the temperature was held at 43°C for the remainder of 15 min). The curd temperature was confirmed to be around 39-40°C. The curd and whey were then incubated at 40°C for another 30 min. After incubation, we removed the curd and whey from the water bath, removed 60 mL of whey from the container, and added 60 mL of sodium chloride solution (0.16g/mL) (salting). The box with curd and whey was returned to the water bath after salting and incubated at 41°C for 20 min. After incubation, the whey was drained from the curd using a sterile cheese cloth and allowed to flow off the curd by gravity. The curd was then cut into small pieces, weighed into 5 g portions and molded into individual wells of a sterile 12-well tissue culture plate. After all the curd was weighed and poured into individual wells of the 12-well tissue culture plate, each mini cheese was pressed by placing a sterile cap of a 15-ml conical tube (topside down) on top of each well. The lid of the 12-well plate was placed on the plate and each plate containing the mini cheeses was flipped upside down to allow effecting draining of any additional whey into a tray. A 0.5 kg weight was placed on top of the 12-well plates containing the mini cheeses, the plates were placed in a secondary container with locked lid and the mini cheeses were pressed for 16 h at 4°C. The 15-ml conical tube caps were removed after pressing, excess whey was drained and the 12-well plates containing the mini cheeses were aged at 4°C.

### Commercial raw milk cheese samples

As part of routine field investigations, the U.S. Food and Drug Administration (FDA) identified commercial raw milk cheeses that were suspected to be HPAI H5N1 virus positive, as the raw milk and raw cheese dairy tested positive for HPAI H5N1 virus. These were cheddar type raw milk cheeses molded in 18 kg blocks that were at day 24 of aging. A total of four 1 kg cheese samples were collected by FDA from the 18 kg cheese blocks and submitted to our laboratory for testing and aging. Upon arrival, 1 g samples were collected from each field cheese block (day 24) and processed for virological assessments as described below. Aging was performed at 4°C as described for the mini cheeses above.

### Sample collection and processing

To assess the stability of HPAI H5N1 virus during the cheese making process and aging, multiple samples were collected from the raw milk mini cheeses and the commercial raw milk cheese studies. During the mini cheese making process, milk samples (1 mL) were collected immediately after HPAI-spiking, after pH adjustment, and after heating at 34°C for 1 h. Additionally, whey samples were collected at different steps of coagulation and whey separation, including: after coagulum formation and incubation at 35°C for 65 min (whey 1), after heating the curd at 40°C for 30 min (whey 2) and after salting of the curd by addition of sodium chloride solution and heating at 40°C for 20 min (whey 3). Additionally, a 1 g sample of the curd was collected after draining of the whey was completed. All samples were stored at -80°C for further virological assessments (**Extended Data Fig. 1**).

To assess HPAI H5N1 stability during raw milk cheese aging, 1 g samples of mini cheeses or of the commercial raw milk cheeses were collected throughout the 60-day aging period recommended by FDA for raw milk cheeses. Samples from the mini cheeses were collected daily from days 1 to 7 and then on days 14, 21, 28, 35, 42, 49, 56 and 60 of aging. Samples (4 x 1 g) from each of the commercial raw milk cheeses were collected twice per week (days 29, 32, 35, 38, 42, 45, 48, 52, 56 and 60) during the 60-day aging period. One 1 g-sample of each cheese was transferred to a sterile 7 oz whirl-pak homogenizer blender filter bag and homogenized with 10 mL sterile phosphate buffered saline (PBS, 10% w/v suspension). The cheese suspension was clarified by centrifuging at 3000 x g for 10 min in a refrigerated centrifuge, and 1 mL aliquots of the cheese homogenate was collected in sterile screw-capped vial and stored at -80°C for virological assessments.

### Monitoring the raw milk and raw milk cheese pH

The pH of raw milk used to produce the mini cheeses and following pH adjustment for cheese production was measured by using a pH meter (HI2002-01 edge® Dedicated pH, Scientific Equipment Source, ON, Canada), and the pH of the mini cheeses and of the commercial raw milk cheeses was measured using a solid pH probe and the same pH meter (HI14140 - Digital flat tip pH electrode for Edge, Scientific Equipment Source, ON, Canada). The pH measurements for mini cheeses were taken on days 0, 7, 14, 21, 28, 35, 42, 49, 56 and 60 of aging while for commercial raw milk cheeses on days 24, 29, 32, 35, 38, 42, 45, 49, 52, 56 and 60.

### Measuring water activity of cheese

The water activity (A_w_) in the mini cheeses and the commercial raw milk cheese was measured using the Aqualab 4TE Water Activity Meter (Aqualab, Australia). Briefly, 3-4 g of each cheese were chopped or minced into small pieces and transferred to Aqualab Sample Cup Bottoms (Aqualab, Australia). The cup was then transferred to the Water Activity Meter and A_w_ was determined. The A_w_ measurements were taken on the same days of sample collection and pH measurements as described above.

### RNA extraction and RT-PCR

Viral RNA from milk, cheese homogenates, or allantoic fluid (200 µl) from embryonated chicken eggs inoculated with the cheese samples was extracted using the IndiMag Pathogen kit (INDICAL Bioscience) on the IndiMag 48s automated nucleic acid extractor (INDICAL Bioscience, Leipzig, Germany) following the manufacturer’s instructions. Real time reverse transcriptase PCR (rRT-PCR) was performed using the Path-ID™ Multiplex One-Step RT-PCR Kit (Thermo Fisher, Waltham, MA, USA) and primers and probes targeting the M gene under the following conditions: 15 min at 48°C, 10 min at 95 °C, 40 cycles of 15 s at 95 °C for denaturation and 60 s at 60 °C. A standard curve was prepared using RNA extracted from HPAI H5N1 TX2/24 spiked milk samples. Serial 10-fold dilutions of the stock virus (2x10^7^ log TCID_50_/mL) were prepared in raw milk samples for RNA extraction followed by RT-PCR as described above. The Ct values were used to estimate the viral RNA copy number in the tested samples using the relative quantification method.

### Virus titration in embryonated chicken eggs (EID_50_)

For virus titration in eggs, 10-day-old embryonated chicken eggs were used. Serial 10-fold dilutions of milk, whey or cheese homogenate samples were prepared in phosphate buffered saline (PBS) supplemented with antibiotic-antimycotic (anti-anti 100X, Thermo Fisher Scientific, USA). The embryonated chicken eggs were candled to mark the air sac on the shell, sanitized with 70% ethanol and a hole was drilled in the eggshell. One hundred µL of the diluted milk or homogenized cheese samples were injected in triplicate into the allantoic cavity route and sealed with glue. Eggs were candled daily for four days, and dead embryos were chilled overnight before collection of allantoic fluid. After 4 days of inoculation, all eggs were chilled for 24 hours, and allantoic fluid was collected. All allantoic fluid samples were tested by hemagglutination assay (HA) using 0.5% chicken RBC. Finally, the 50% egg infectious dose (EID_50_) was calculated using the Reed and Muench method.

The presence of influenza A virus in the allantoic fluid of ECE inoculated with select cheese homogenate samples (day 60 of aging for mini cheeses or days 24, 42, 45 and 60 of aging for commercial raw milk cheeses) was confirmed by rRT-PCR as described above. Additionally, influenza A whole genome sequencing was performed on commercial raw milk cheese samples and allantoic fluids collected on days 24 and 60 of aging to confirm the presence of H5N1 virus.

### Hemagglutination (HA) assay

For the HA assay, 100 µL allantoic fluid was taken into the wells of 1^st^ column of a 96-well U-bottom plate. 50 µL of PBS was added to the wells of 2^nd^ and 3^rd^ columns. Two 2-fold dilutions (1:2 to 1:8) of the allantoic fluids were prepared in PBS. Then, 50 µL of 0.5% chicken red blood cells (RBC) were added to each well and incubated at room temperature for 30 min. Lack of RBC button formation indicates positive HA reaction.

### Next generation sequencing

The whole genome sequence of influenza A present in the commercial raw milk cheese samples was determined using targeted influenza A sequencing at the Virology Laboratory at the Cornell Animal Health Diagnostic Center as previously described^1^. Sequence and phylogenetic analyses were performed as previously described^1^.

### Determination of *D*-values

The thermal inactivation kinetics of HPAI H5N1 in raw milk cheese were calculated based on the decimal reduction time (*D*-value). The *D*-value is defined as the time required at a specific temperature to achieve a one-logarithm reduction in viral titer. This was determined by plotting the logarithm (base 10) of the infectious viral titers against time for cheese made with milk at pH 6.6 and 5.8. The *D*-value was then calculated as the negative inverse of the slope of the resulting plot, with the line of best fit for the survivor curves established through regression analysis.

### pH dependent infectivity inhibition of HPAI H5N1 virus in raw milk

Cal-1 cells were seeded in 24-well plate (2.5x10^5^ cells/mL) and incubated at 37°C for 24 h. Raw normal bulk tank milk was spiked with the recombinant HPAI H5N1 TX2/24-NLuc virus to obtain a target titer of about 10^7^ EID_50_ or 10^6^ TCID_50_ per mL of milk. The pH of the raw bulk tank milk was left unadjusted at pH 6.78 or adjusted to pH 6.0, 5.5, 5.0 and 4.5 and incubated at 4°C for 1 hour. Cal-1 cells were washed with sterile PBS and 200 µl of the milk samples were added to each well. Wells inoculated with raw normal milk were kept as a control. The plate was transferred to refrigerator and incubated for 1 h (virus adsorption) at 4°C. After incubation, cells were washed twice with sterile PBS and 500 µL of complete growth medium (MEM 10% FBS) was added to each well. The plate was then incubated at 37°C for 4 h. After incubation, cell supernatant was aspirated, 50 µL of 1X Passive Lysis Buffer (Promega) was added to each well and incubated at room temperature for 10 min. After 1 cycle of freeze-thaw, 10 µL of cell lysate was transferred to each well of a luminometer plate to which 50 µL of luciferase assay reagent (Nano-Glo, Promega) was added. Luminescence was measured using a luminometer plate reader (BioTek Synergy LX Multimode Reader).

### Statistical analysis

Graphs were prepared using GraphPad Prism 10 software.

